# Amphiphilic protein surfactants reduce the interfacial tension of biomolecular condensates

**DOI:** 10.1101/2025.06.19.660548

**Authors:** Bruna Favetta, Huan Wang, Zheng Shi, Benjamin S. Schuster

## Abstract

Biomolecular condensates are protein-dense regions in cells that often arise from liquid-liquid phase separation. Interfacial tension is a key determinant of biomolecular condensate behavior, influencing condensate size and interactions with intracellular structures. Certain proteins and RNAs are known to selectively localize to the interface of condensates, where they can regulate condensate function in cells. Previously, we designed amphiphilic proteins that preferentially adsorb to the surface of condensates. These proteins contain one phase-separating domain (RGG) and one non-phase-separating domain (MBP or GST). Here, we demonstrate through direct quantification that these amphiphilic proteins act as surfactants, reducing the interfacial tension of RGG-RGG condensates from ∼260 µN/m to ∼100 µN/m in a concentration-dependent manner. Notably, the GST-based surfactant protein exhibits a 10-fold greater efficacy in lowering interfacial tension compared to the MBP-based surfactant. We show that this increased efficacy is due to its higher surface density, driven by GST’s ability to oligomerize. We also show that these surfactant proteins slow droplet fusion and reduce average droplet size, as would be expected of a typical surfactant. Our findings quantitatively show how surfactant proteins can play a critical role in regulating the behavior of biomolecular condensates by modulating their interfacial tension.

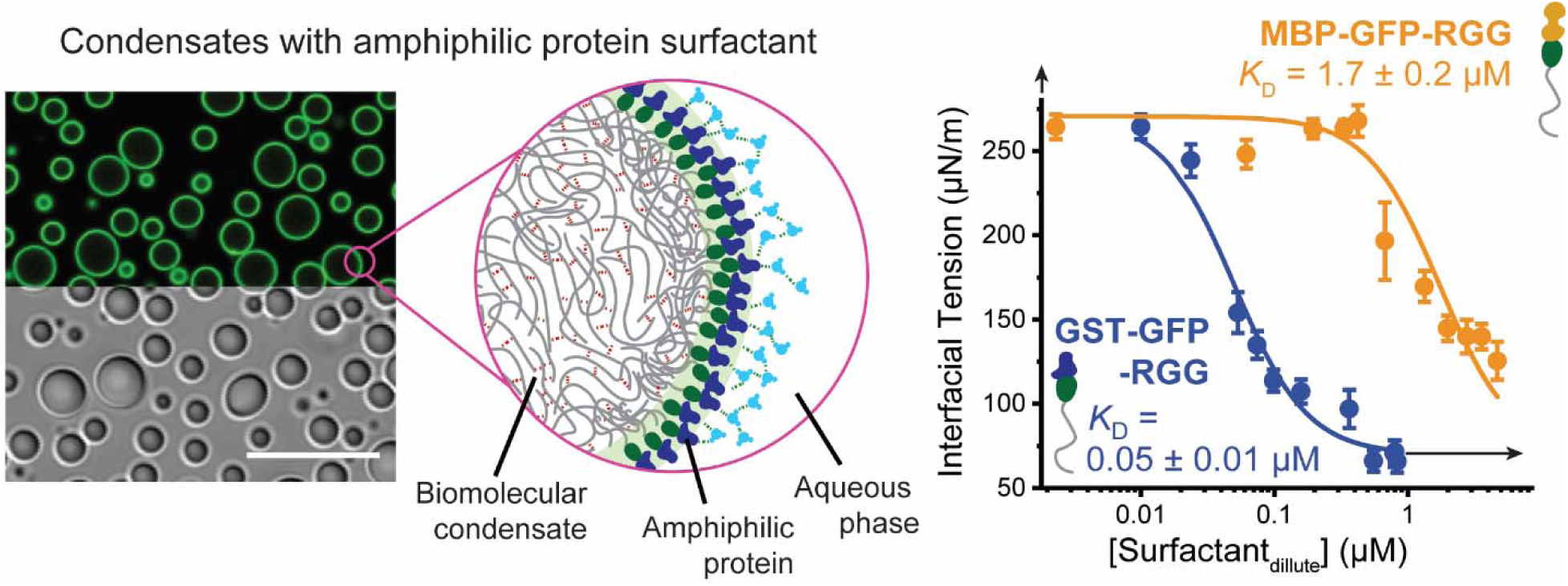

## Introduction

Biomolecular condensates are membraneless organelles that often arise from liquid-liquid phase separation, organizing key cellular processes such as transcription, signal transduction, and stress response^1,2^. The physical properties of these condensates, including their interfacial tension, play a critical role in determining their size, miscibility, dynamics, and interactions with other cellular structures^3,4^. For example, the different interfacial tensions of nucleolar condensates determine the multiphase architecture of the nucleolus^5^. The interfacial tension of DNA-protein condensates can affect the rate of gene expression^6^. Also, the degree of condensate wetting, partly determined by interfacial tension of the condensate, regulates the process of autophagosome formation^7^.

The interfacial tension of biomolecular condensates arises from the balance of intermolecular interactions between proteins and their surrounding environment^3^. Inside the condensate, attractive interactions, such as hydrogen bonding, hydrophobic interactions, and electrostatic forces between amino acids promote cohesion among proteins^8^. At the interface with the surrounding medium, these interactions are reduced, leading to an energetic penalty for exposing certain residues or domains to the surrounding environment. This energetic penalty creates a tension at the interface, acting to minimize the interfacial area of condensates^9^. Because proteins are macromolecules,^10^ and because protein-protein interactions in condensates are often polar and thus similar in nature to those in the aqueous dilute phase^8^, the interfacial tension of condensates is low compared to that of more commonly studied emulsions, such as oil-water^11^. Still, due to this interfacial tension, condensates coarsen over time, either through Ostwald ripening or droplet coalescence^12,13^. The distribution of condensate size is thought to be regulated in cells^14–16^ and has been shown to be important for condensate function^17,18^.

One mechanism that has been proposed to be used by cells to control condensate size is surfactant-like proteins^17,19^. Surfactants are molecules that lower interfacial tension and kinetically stabilize emulsions^20^. Several proteins that interact with condensates in cells have been suggested to have surfactant-like features. During mitosis, the proliferation marker protein Ki-67 prevents chromosomes from merging into a single chromatin mass^21^. However, whether Ki-67 affects interfacial tension has not been measured. In a second example, the protein NO145 preferentially partitions to the interface of *X. laevis* nucleoli and may prevent fusion of nucleoli^14^. In a third example, MLX, a transcription regulator, preferentially locates to the surface of condensates composed of the transcription factor TFEB in vitro and lowers the interfacial tension of TFEB condensates by around 2.5 times, thus decreasing their affinity for a specific DNA motif^22^. However, the mechanism of regulation of interfacial tension was not explored. In general, factors that determine a condensate’s interfacial tension are poorly understood. Consequently, developing model surfactant proteins and examining how they influence condensates can provide valuable insights into the underlying mechanisms of condensate interfacial tension regulation in cells.

In previous work, we designed amphiphilic proteins containing an intrinsically disordered region (IDR) fused to folded domains^19^. We utilized the soluble folded domain glutathione S-transferase (GST) or maltose binding protein (MBP), which we fused to the intrinsically disordered RGG domain of P granule protein LAF-1. This RGG domain has been previously shown to be necessary and sufficient for LAF-1 phase separation^23^. RGG is a 168-residue domain enriched in Arg, Gly, Tyr, and uncharged polar residues^24^. Two amphiphilic protein constructs we designed previously, and further characterize here, are MBP-GFP-RGG and GST-GFP-RGG^19^. At low concentrations, these amphiphilic proteins form a film that coats the surface of a condensate composed of RGG-RGG protein (a tandem repeat of the RGG domain). This can be understood as follows: In the amphiphilic fusion proteins, the RGG domain is “condensate-philic”, thus preferring to interact with the RGG-RGG condensate. The folded MBP or GST domains are “condensate-phobic”, thus preferring to interact with the dilute phase. We included GFP to visualize the localization of the amphiphilic proteins.

Here, we directly measure the effect of these amphiphilic proteins on the interfacial tension of RGG-RGG condensates. Using micropipette aspiration, we find that both amphiphilic proteins reduce condensate interfacial tension from approximately 260 µN/m to approximately 100 µN/m in a concentration-dependent manner. However, the GST-based surfactant is ∼10 times more efficient compared to the MBP-based surfactant. Furthermore, we show that the efficiency of interfacial tension reduction depends on the ability of amphiphilic proteins to form oligomers. The effect of amphiphilic proteins on condensate interfacial tension can be rationalized by a model in which increased amphiphilic protein density at the condensate surface directly reduces the interfacial tension. Consistent with their ability to reduce condensate interfacial tension, we observe that amphiphilic proteins slow the fusion and reduce the size of condensates. Collectively, our results establish a quantitative relationship between the presence of surfactant proteins and condensate interfacial tension, shedding light on how cells may regulate several aspects of condensate functions.

## Experimental Section

### Cloning

All genes of interest were cloned into pET vectors in frame with C-terminal 6x-His tags. RGG-RGG was cloned as previously described^25^. Amphiphilic proteins were cloned as described previously^19^. The RGG domain used here is the N-terminal IDR (residues 1-168) of *C. elegans* P granule protein LAF-1^26^. Gene sequences were verified by Sanger sequencing (Azenta) and are available in SI Appendix Note 1.

### Protein expression and purification

For bacterial expression, plasmids were transformed into BL21(DE3) competent *E. coli* (New England BioLabs). Colonies picked from fresh plates were grown for 8 h at 37 °C in 1 mL LB while shaking at 250 rpm. This starter culture was then used to inoculate 0.5 L cultures. For GST-MBP-RGG and MBP-GFP-RGG, cultures were grown in 2 L baffled flasks in Terrific Broth medium (Fisher Scientific) supplemented with 4 g/L glycerol while shaking at 250 rpm. The flasks were shaken at 37°C until the OD600 reached approximately 1, at which time expression was induced with 500 µM isopropyl β-D-1-thiogalactopyranoside (IPTG) and the temperature was reduced to 18°C for overnight expression. For RGG-RGG, cultures were grown in 2 L baffled flasks in Autoinduction Medium (Formedium) supplemented with 4 g/L glycerol at 37 °C overnight while shaking at 250 rpm. The pET vectors used contained a kanamycin resistance gene; kanamycin was used at concentrations of 50 μg/mL in cultures. After overnight expression at 18 °C or 37 °C, bacterial cells were pelleted by centrifugation at 4100 x g at 4 °C. Pellets were resuspended in lysis buffer (1 M NaCl, 20 mM Tris, 20 mM imidazole, Roche EDTA-free protease inhibitor, pH 7.5) and lysed by sonication. Lysate was clarified by centrifugation at 25000 x g for 30 minutes at 25 °C. The clarified lysate was then filtered with a 0.22 µm filter. Lysis was conducted on ice, but other steps were conducted at room temperature to prevent phase separation.

Proteins were purified using an AKTA Pure FPLC with 1 mL nickel-charged HisTrap columns (Cytiva) for immobilized metal affinity chromatography of the His-tagged proteins. After injecting proteins onto the column, the column was washed with 500 mM NaCl, 20 mM Tris, 20 mM imidazole, pH 7.5. Proteins were eluted with a linear gradient of imidazole up to 500 mM NaCl, 20 mM Tris, 500 mM imidazole, pH 7.5. Proteins were dialyzed overnight using 7 kDa MWCO membranes (Slide-A-Lyzer G2, Thermo Fisher) into physiological buffer (150 mM NaCl, 20 mM Tris, pH 7.5), with the exception that GST-GFP-RGG was dialyzed into high salt buffer (500 mM NaCl, 20 mM Tris, pH 7.5). Proteins were dialyzed at room temperature (20 °C) except for RGG-RGG, which was dialyzed at 42 °C to inhibit phase separation, because the phase-separated protein bound irreversibly to the dialysis membrane. Proteins were snap frozen in liquid nitrogen in single-use aliquots and stored at −80 °C.

### Confocal Microscopy

Protein samples were prepared as follows: RGG-RGG protein aliquots were thawed at 42°C, above its phase transition temperature. MBP- and GST-based proteins were thawed at room temperature. Protein concentrations were measured based on their absorbance at 280 nm using a Nanodrop spectrophotometer (ThermoFisher); RGG-RGG was mixed in a 1:1 ratio with 8 M urea to prevent phase separation during concentration measurements.

First, RGG-RGG and buffer were mixed at room temperature. Then, amphiphilic proteins MBP-GFP-RGG or GST-GFP-RGG were added at the desired protein concentrations. The final buffer conditions were 150 mM NaCl, 20 mM Tris, pH 7.5. Protein samples were plated on 16-well glass-bottom dishes (1.5 glass thickness; Grace Bio-Labs) that were coated with 5% Pluronic F-127 (Sigma-Aldrich) for a minimum of 10 minutes to create a hydrophilic surface and prevent condensate wetting the glass. The chambers were washed with water prior to plating the protein samples.

Confocal imaging was performed on a Zeiss Axio Observer 7 inverted microscope equipped with an LSM900 laser scanning confocal module and employing a 63x/1.4 NA plan-apochromatic, oil-immersion objective. GFP was excited to fluoresce with a 488 nm laser. Confocal fluorescence images were captured using GaAsP detectors. Transmitted light images were collected with either the ESID module or an Axiocam 702 sCMOS camera (Zeiss), in both cases using a 0.55 NA condenser.

### Droplet image analysis

Image analysis and data processing were performed in MATLAB R2024b. The fluorescence intensity profile of the condensates with GFP-tagged proteins was measured by using the Circular Hough Transform (imfindcircles function) to identify droplet locations and drawing a line that spanned the droplet diameter plus 25% of a radius length in each direction across the droplets. Condensate size was also calculated using the Circular Hough Transform (imfindcircles function).

### Micropipette Aspiration (MPA)

The micropipette aspiration (MPA) experiments were carried out on a Ti2-A inverted fluorescence microscope (Nikon, Japan), with a 60x water objective (NA 1.2), a Hamamatsu camera (ORCA-Fusion, Flash4.0 V3, Hamamatsu), equipped with a motorized stage and two motorized 4-axes micromanipulators (PatchPro-5000, Scientifica) and a multi-trap optical tweezers (Tweez305, Aresis, Slovenia) according to the protocol we reported previously^11,27^ with minor modifications.

Micropipettes were made by pulling glass capillaries using a pipette puller (PUL-1000, World Precision Instruments). The pipette tip was then cut to achieve an opening diameter ranging from 2 to 5 μm. Subsequently, the pipette was bent to an angle of approximately 40° using a microforge (DMF1000, World Precision Instruments) so that the tip of the micropipette would be parallel to the imaging plane. The micropipette was filled with the same buffer as used in other microscopy experiments (150 mM NaCl, 20 mM Tris, pH 7.5) using a MICROFIL needle (World Precision Instruments). The filled micropipette was then mounted onto a micromanipulator. The rear end of the pipette was connected to an automatic pressure controller (Flow-EZ, Fluigent; Pressure resolution 1 Pa). The MPA experiments were conducted in glass-bottom dishes (D35-20-1.5-N, Cellvis), under brightfield illumination to minimize potential artifacts associated with fluorescence excitation. Fluorescence images were taken after MPA experiments to confirm the presence of the core-shell structure. Typically, optical tweezers-assisted condensate fusion was first carried out to achieve a large (diameter > 5 μm) condensate for accurate MPA measurements. To minimize sample evaporation, 1.5 mL Milli-Q water was added to the edge of the dish (separate from the sample), and the dishes were covered with a thin plastic wrap with a ∼2 mm hole for pipette insertion. Zero pressure of the aspiration pipette was calibrated before each set of experiments by determining the pressure at which small condensates underwent Brownian motion inside the micropipette.

Measurement of condensate viscosity was carried out as described in Wang et al^11^ and analyzed according to the protocol described in Roggeveen et al^27^. Briefly, normalized aspiration length (aspiration length, *L*_p_, over the pipette radius, *R*_p_) was measured over time for each pressure step. For each step, the slope of a linear fitting of (*L*_p_/*R*_p_)^2^ vs. time is equal to the effective shear rate. Then, the slope of aspiration pressure vs. shear rate graph gives 4η, where η is viscosity. In principle, the intercept is the interfacial tension of the condensate. In our case, measurement would need to be taken infinitely slowly to account for equilibration of the surfactant, and so the intercept was not used to measure interfacial tension.

Instead, to more accurately quantify the interfacial tension of condensates, a static tension measurement protocol was used^28,29^. A large condensate was drawn into the micropipette by a small suction pressure (typically ∼ 5 Pa), then the pressure was increased stepwise until the deformation of the condensate interface was equal to the inner radius of the micropipette (SI Appendix, Figure S1). During this process, images were collected at each pressure step. The radii of the condensate protrusion and external condensate droplet were measured for each pressure step using ImageJ (version 2.1). For each step, the interfacial tension, γ, was calculated using^28^:

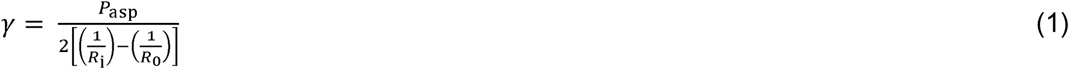

where *R*_i_ is the radius of curvature of the condensate interface inside the pipette, *R*_o_ is the radius of the condensate outside the pipette, and *P*_asp_ is the aspiration pressure used. Interfacial tensions measured from each step were averaged such that all pressure steps received the same weight to obtain the interfacial tension of the condensate.

### Optical Tweezer-Assisted Droplet Fusion

An optical tweezers-assisted fusion assay was carried out and analyzed following the protocol previously described^11^. Two condensates were independently trapped with minimal laser power. The fusion events were captured at 20 Hz. MATLAB R2024b was used to fit the contour of the fusing condensates to an ellipse. The fusion time (r) was found by fitting the relaxation in aspect ratio (AR; defined as the ratio between the long and short axis of the ellipse) over time (*t*) to a stretched exponential decay^30^, according to the following equation:

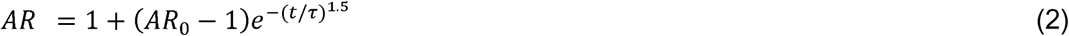

where AR_0_ is the aspect ratio at time 0, defined as the start of the droplet fusion event.

## Results and Discussion

### Amphiphilic Proteins Reduce the Interfacial Tension of RGG-RGG Condensates

In agreement with our previous results, both amphiphilic proteins tested, MBP-GFP-RGG and GST-GFP-RGG, form a film surrounding an RGG-RGG condensate core when 10 µM RGG-RGG is mixed with 1 µM of the amphiphilic protein (Figure 1A-C).

**Figure 1:**
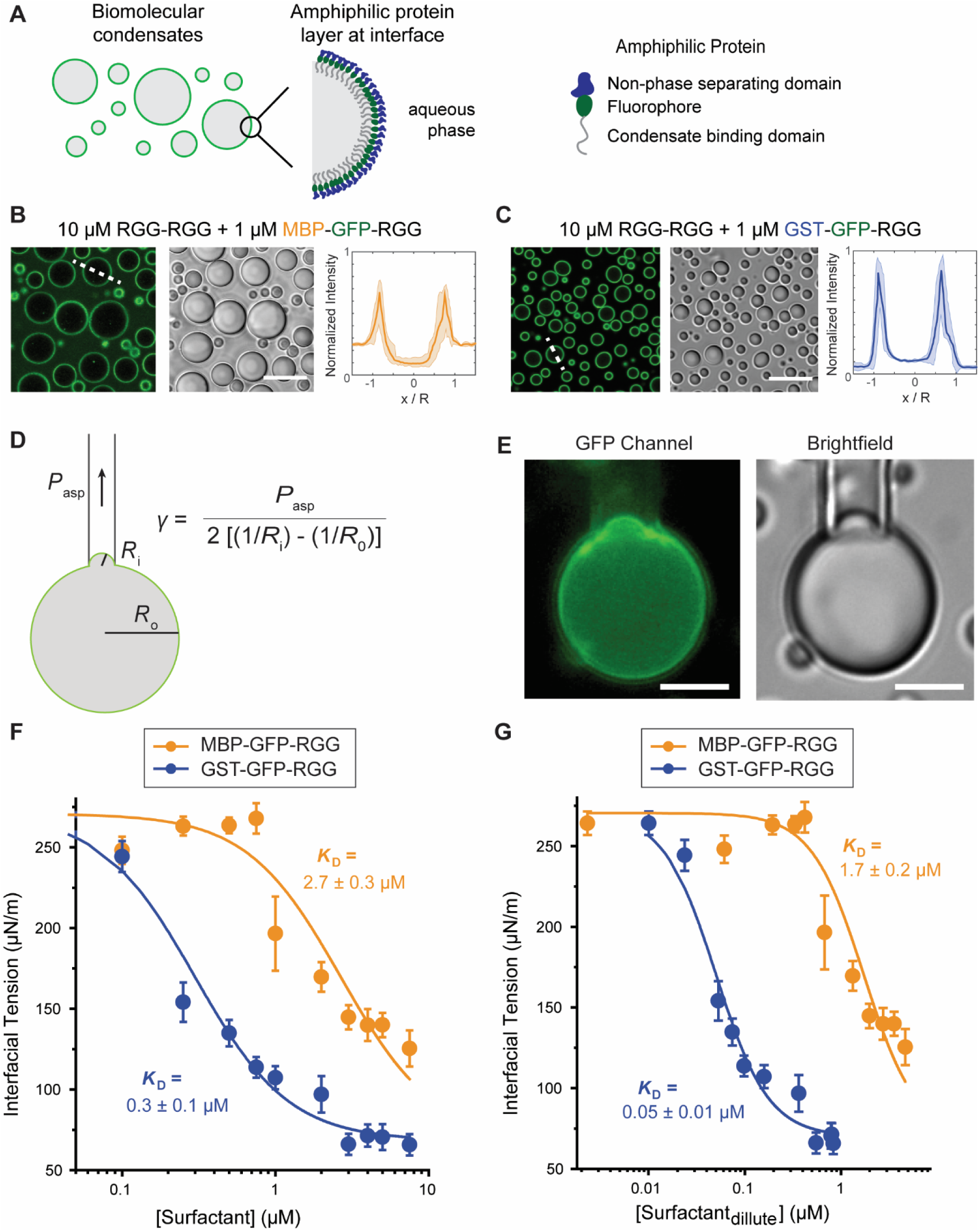
Amphiphilic proteins lower condensate interfacial tension. A) Schematic model of the engineered amphiphilic proteins localized on the surface of an RGG-RGG condensate. B and C) Transmitted light and confocal fluorescence imaging of core-shell condensates formed by mixing 10 μM RGG-RGG and 1 μM MBP-GFP-RGG or GST-GFP-RGG. Fluorescence intensities were quantified as line profiles across individual condensates, normalized by condensate size. Intensities were also divided by 65,535, the dynamic range of the images, such that intensities scale from 0 to 1. N > 50 condensates, from images taken from three independent samples. Shaded area represents 1 standard deviation. (Scale bars, 10 μm). D) Schematic representation of experimental setup for measuring interfacial tension using micropipette aspiration and the corresponding equation. E) Widefield imaging of condensate aspiration experiment, showing partially aspirated condensate retains amphiphilic protein layer at interface (Scale bars, 5 μm). F) Interfacial tension of RGG-RGG condensates measured with increasing concentrations of either amphiphilic protein, MBP-GFP-RGG and GST-GFP-RGG. G) Interfacial tension of RGG-RGG condensates measured against the equilibrium dilute phase concentration of either amphiphilic protein. The lines in F and G represent fitting curves using the Hill equation (Equation 3). Error bars indicate ±1 standard error of the mean.

To measure the interfacial tension of condensates with amphiphilic proteins, we use Micropipette Aspiration (MPA), as described in Figure 1D-E and Methods section *Micropipette Aspiration (MPA)*. We found that both amphiphilic proteins behave as surfactants and reduce the interfacial tension of RGG-RGG condensates in a surfactant concentration-dependent manner (Figure 1F), while minimally impacting the viscosity of the condensates (SI Appendix, Figure S2). For MBP-GFP-RGG, at concentrations up to 0.75 µM, we observe a plateau in the interfacial tension measurements at ∼260 µN/m. As the concentration of MBP-GFP-RGG continues to increase, the interfacial tension of condensates significantly decreases and reaches ∼125 µN/m at 7.5 µM amphiphilic protein. For GST-GFP-RGG, a significant decrease in interfacial tension is observed with much smaller concentrations of the amphiphilic protein, as low as 0.25 µM. With increasing GST-GFP-RGG concentrations, we observe a reduction in interfacial tension until reaching a plateau at ∼65 µN/m.

We next sought to quantitatively define the interfacial effects of each surfactant protein.

Given the sigmoidal shape of the interfacial tension (y) vs. surfactant concentration (C) curves, the traditional Szyszkowski–Langmuir equation cannot fully capture the trend of the data (SI Appendix, Figure S3). We therefore fit our results to the Hill equation^31^, assuming that the density of surfactants adsorbed to the interface of condensates linearly reduces the interfacial tension (SI Appendix Note 2):

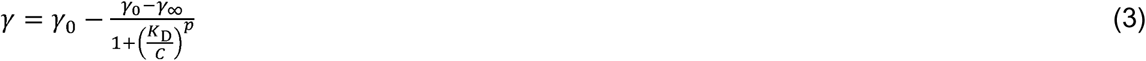

Here, *C* is the surfactant concentration, γ_0_ and γ_∞_ are the initial and final interfacial tension, and *K*_D_ and *p* are the dissociation constant and cooperativity coefficient of the interfacial adsorption of the surfactants, respectively. To minimize fitting uncertainty, we carried out a global fit to both the MBP-GFP-RGG and the GST-GFP-RGG data with γ_0_, γ_∞_, and *p* as shared parameters and only *K_D_* as a free fitting parameter. We find that for MBP-GFP-RGG, *K*_D_ = 2.7 ± 0.3 μM, whereas for GST-GFP-RGG, *K*_D_ = 0.3 ± 0.1 μM. Therefore, GST-GFP-RGG is 9-times more efficient than MBP-GFP-RGG in reducing the interfacial tension of RGG-RGG condensates (Figure 1F). We also found *p* = 1.6 ± 0.3, suggesting a cooperative binding mechanism for surfactant proteins, perhaps because binding of surfactant molecules induces ordering of RGG-RGG molecules at the interface, which promotes the adsorption of additional surfactant molecules^32^.

Interestingly, our results indicate a correlation between the brightness of fluorescence at the condensate interface and the efficiency of the amphiphilic protein in reducing interfacial tension. As noted above, we can observe the degree of protein adsorption to the condensate surface through confocal fluorescence imaging. We measured the degree of partitioning of MBP-GFP-RGG and GST-GFP-RGG in the dilute phase, interface, and condensed phase (Figure 1B-C). Fluorescence intensities were quantified as line profiles across individual condensates, normalized by condensate size. Intensities were also divided by 65,535, the dynamic range of the 16-bit images, such that intensities scale from 0 to 1. At 1 μM amphiphilic protein, the average partitioning for MBP-GFP-RGG is 0.26 ± 0.02 : 0.70 ± 0.09 : 0.11 ± 0.03 in the dilute phase, interface and condensed phase, respectively. For GST-GFP-RGG, we found an average partitioning of 0.04 ± 0.01 : 0.89 ± 0.02 : 0.14 ± 0.03, respectively. Therefore, the GST-based amphiphilic protein prefers the interface more than the MBP-based amphiphilic protein. The increased partitioning to the condensate surface by the GST-based amphiphilic protein likely contributes to its greater strength as a surfactant, an observation that we further explore in Figure 2.

**Figure 2:**
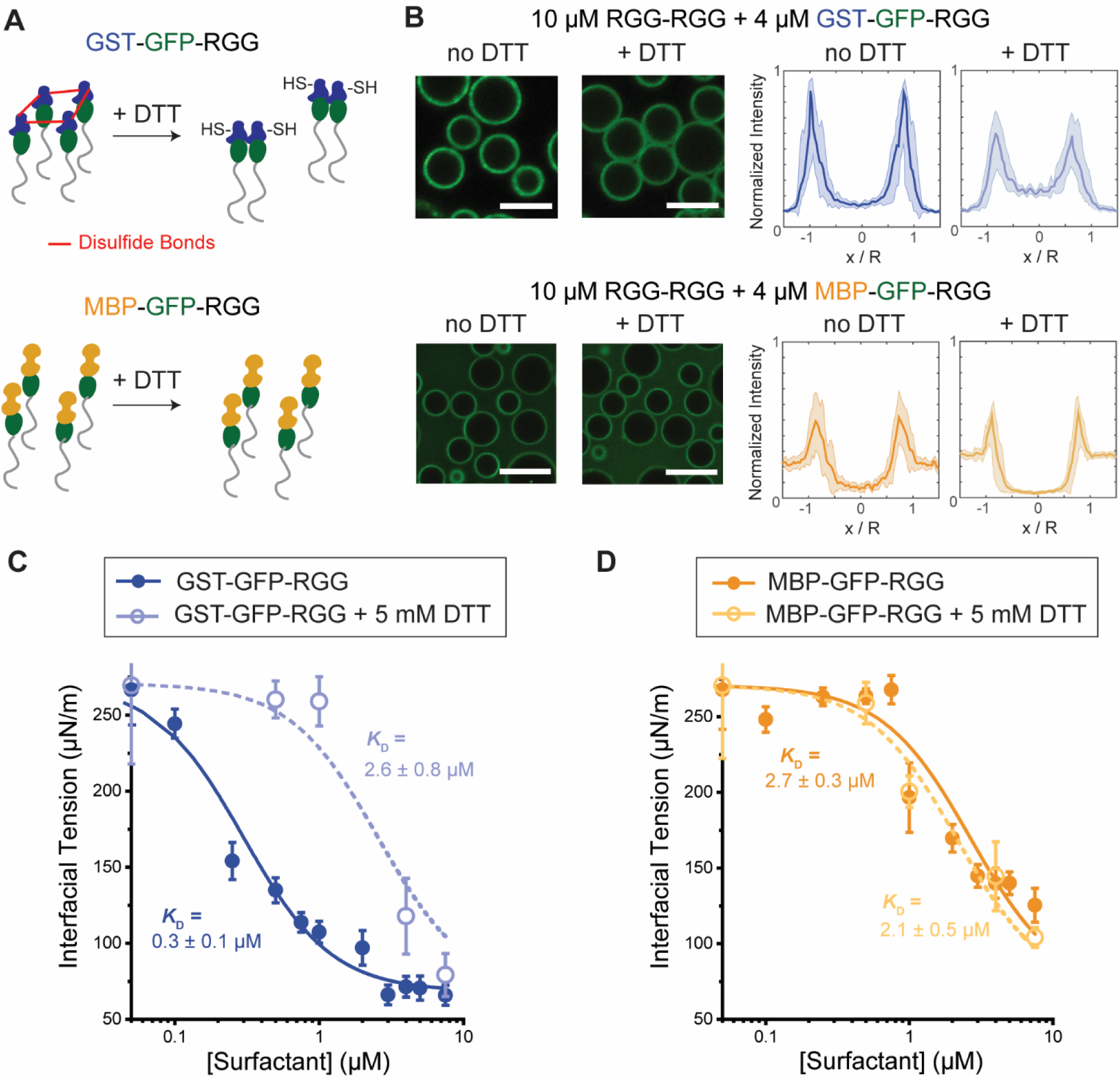
Modulation of surfactant interfacial behavior using reducing agent DTT. A) Schematic depicting that DTT inhibits disulfide bond formation between cysteines in the GST domains, while not affecting MBP. B) Confocal fluorescence imaging of the condensates shows adding 5 mM DTT lowered the partitioning of GST-GFP-RGG, with no effect on MBP-GFP-RGG (Scale bars, 5 μm). Fluorescence intensities were quantified as line profiles across individual condensates, normalized by condensate size. Intensities were also divided by 65,535, the dynamic range of the images, such that intensities scale from 0 to 1. N > 20 condensates, from images taken from two independent samples. Shaded area represents 1 standard deviation. C and D) The effect of GST-GFP-RGG (C) and MBP-GFP-RGG (D) on condensate interfacial tension with and without the addition of DTT. The lines represent fitted curves to the Hill equation. Error bars indicate 1 standard error of the mean.

The partitioning analysis above also indicates that there is a difference between the equilibrium dilute phase concentrations of samples with GST-GFP-RGG and MBP-GFP-RGG. This suggests that there may be a difference between the equilibrium dilute phase concentrations and the total concentration initially added of the amphiphilic proteins. To account for this effect, we generated a standard curve relating GFP fluorescence intensity to the molar concentration of each amphiphilic protein, using the same buffer solutions as in our experiments (SI Appendix, Figure S4). We then used the standard curve along with our confocal images of condensates to determine the equilibrium concentration of amphiphilic proteins in the dilute phase for each concentration tested, allowing us to re-plot the interfacial tensions as a function of equilibrium surfactant concentrations in the dilute phase (Figure 1G). Using this approach, the difference in *K*_D_ between MBP-GFP-RGG and GST-GFP-RGG increases from 9 times to 34 times – further supporting our finding that the GST-GFP-RGG amphiphile is more efficient at reducing interfacial tension.

### Oligomerization Increases Surfactant Strength

Next, we investigated the cause of the difference in surfactant strength between the two amphiphilic proteins. Unlike MBP, which is monomeric and lacks cysteines, GST is dimeric and additionally has a high tendency to oligomerize due to its four cysteine residues (per monomer), three of which are highly solvent exposed and can easily form disulfide bonds^33^. These disulfide bonds can be reduced by dithiothreitol (DTT) (Figure 2A). Therefore, we tested the effect of DTT on the partitioning of surfactant proteins and their ability to decrease interfacial tension. After adding 5 mM DTT, the partitioning of GST-GFP-RGG to the interface of RGG-RGG condensates was reduced, compared to no observable change for the partitioning of MBP-GFP-RGG (Figure 2B). For example, in samples with 4 μM GST-GFP-RGG, the partitioning of amphiphilic protein shifted from 0.09 ± 0.02 : 0.90 ± 0.08 : 0.16 ± 0.04 in the dilute phase, interface and condensed phase, respectively, without DTT, to 0.10 ± 0.03 : 0.58 ± 0.08 : 0.24 ± 0.06, with DTT, which represents a 35% reduction in partitioning to the interface.

To assess whether DTT affects the amphiphilic proteins’ ability to decrease condensate interfacial tension, we repeated our MPA measurements with the addition of 5 mM DTT (Figure 2C - D). Again, we fit our data to the Hill equation (Equation 3) and carried out a global fit to both the MBP-GFP-RGG and the GST-GFP-RGG data with γ_0_, γ_∞_ and *p* as shared parameters and only *K_D_* as a free fitting parameter. We found that DTT has a negligible effect on the interfacial tension of RGG-RGG condensates with MBP-GFP-RGG surfactant (*K_D_* = 2.7 ± 0.3 µM without DTT and 2.1 ± 0.5 µM with DTT). However, DTT strongly impairs the surfactant activity of GST-GFP-RGG, with *K_D_* increasing upon adding DTT from 0.3 ± 0.1 µM to 2.6 ± 0.8 µM, which is comparable to that of MBP-GFP-RGG. Thus, our findings suggest that the ability of GST to oligomerize and form disulfide bonds drives its higher partitioning to the condensate surface and increases its strength as a surfactant. In prior work, we showed that the strength of interaction between “condensate-phobic” domains partially determines the degree of partitioning of an amphiphilic protein to the surface of a condensate^19^. Here, we found that the greater partitioning of GST-based amphiphiles to the surface of condensates also correlates to a greater reduction in interfacial tension. Our findings also suggest that modulation of the redox environment is one mechanism that can be used to regulate the strength of a protein surfactant, which we speculate may be relevant in cells.

### Amphiphilic Proteins Slow Droplet Fusion and Reduce Condensate Size

Droplet coalescence is driven by interfacial tension. We therefore asked whether surfactants reduce condensate fusion time. Here, we utilized an optical tweezer-assisted droplet fusion assay to quantify fusion time and the inverse capillary velocity of condensates at various concentrations of amphiphilic proteins. As shown in Figure 3A and quantified in Figure 3B, we observed that addition of either amphiphilic protein increased condensate fusion time. We fit the merging condensates to a stretched exponential equation to quantify the change in aspect ratio over time^30^. Two ∼4 µm radius RGG-RGG condensates fused within 0.4 seconds, whereas condensates of similar sizes in samples with 5 µM MBP-GFP-RGG fused within 0.8 seconds, and those with 5 µM of GST-GFP-RGG fused in 2 seconds.

**Figure 3:**
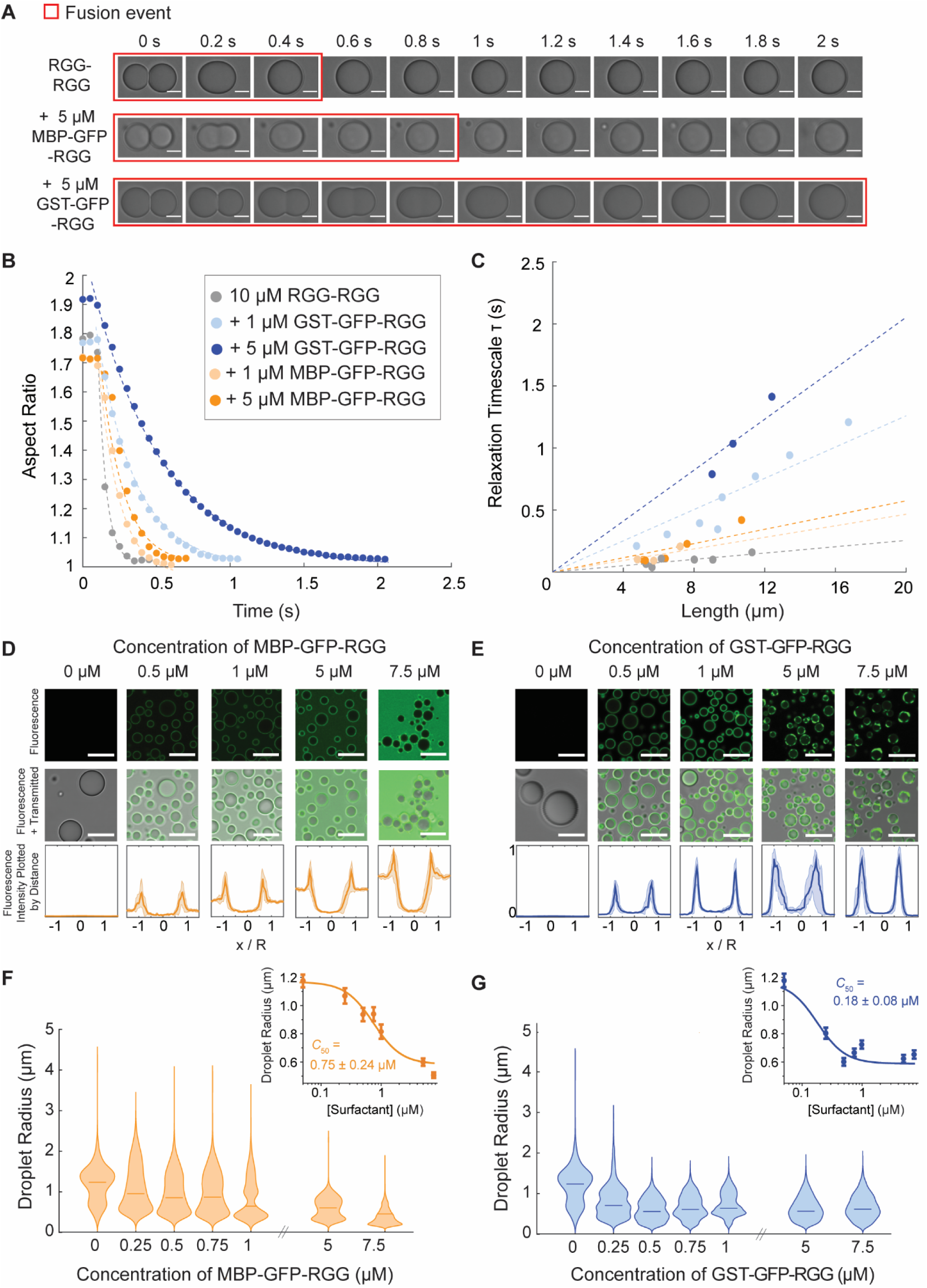
Amphiphilic proteins slow droplet fusion and also reduce the average size of condensates. A) Time-lapse brightfield images of optical tweezer-assisted fusion of condensates, with and without the amphiphilic proteins. Red boxes represent the fusion events, where the aspect ratio changes from ∼2 to 1. B) Relaxation of condensate aspect ratios over time with and without the addition of either amphiphilic protein, comparing similarly sized (∼4 μm radii) condensates. Increasing the concentration of amphiphilic proteins slows the relaxation of the aspect ratio. C) Relaxation time scale of droplet fusion, plotted against droplet length (the geometric mean of condensate diameter before fusion). The dashed lines are linear fits for each condition. D and E) Transmitted light and confocal fluorescence imaging of the condensates formed by mixing RGG-RGG (10 μM) with different concentrations of MBP-GFP-RGG (D) and GST-GFP-RGG (E). Droplet radii become smaller as amphiphilic protein concentration increases. Fluorescence intensities were quantified as line profiles across individual condensates, normalized by condensate size. Intensities were also divided by 65,535, the dynamic range of the images, such that intensities scale from 0 to 1. N > 50 condensates, from images taken from three independent samples. Shaded area represents 1 standard deviation. Scale bars, 5 μm. F and G) Violin plots depicting shift in size of condensates with increasing concentrations of MBP-GFP-RGG (F) and GST-GFP-RGG (G). Bars indicate the median droplet size in each condition. Inset, the relationship between condensate size and the concentration of either amphiphilic protein, MBP-GFP-RGG and GST-GFP-RGG fit to a sigmoidal equation. C_50_, the concentration required to reduce average condensate radius by half, is noted. Error bars indicate 1 standard error of the mean.

Next, we extracted relaxation timescales and plotted those against condensate radius for multiple fusion events (Figure 3B-C) to obtain the inverse capillary velocity, η/γ:

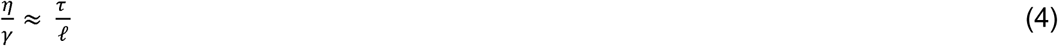

where η is the viscosity, γ is the interfacial tension, r is the relaxation timescale, and ℓ is the condensate length determined as the geometric mean of the condensate diameters prior to fusion^30,34^. Our results indicate that increasing concentrations of amphiphilic protein increases the slope and therefore the inverse capillary velocity of samples. For RGG-RGG, we find an inverse capillary velocity of 0.013 s/µm, compared to 0.023 and 0.030 s/µm for samples with 1 and 5 µM MBP-GFP-RGG, versus 0.064 and 0.104 s/µm for samples with 1 and 5 µM GST-GFP-RGG. We can also calculate inverse capillary velocity from our micropipette aspiration data, from which we determined viscosity (SI Appendix S2) and interfacial tension (Figure 1).

From our micropipette aspiration data, we obtain inverse capillary velocities of 0.014 s/µm for RGG-RGG, 0.025 for samples with 1 µM MBP-GFP-RGG and 0.047 for samples with 1 µM GST-GFP-RGG. Comparing the results, we find agreement between the inverse capillary velocities calculated from the fusion data compared to that from the micropipette aspiration data (SI Appendix, Table 1). Therefore, using the concept of inverse capillary velocity, we can connect the reduction in droplet fusion time to the reduction in interfacial tension in samples with amphiphilic proteins.

Interestingly, we also noted cases with > 5 µM of amphiphilic protein concentration where we could not induce droplet fusion with our optical tweezers (SI Appendix, Movie 1). This result suggests that the amphiphilic protein film at the surface of condensates acts as a barrier that significantly hinders condensate fusion beyond the expected effect of interfacial tension. Therefore, we next tested whether the amphiphilic proteins affect the condensate size distribution. For this, we measured droplet sizes in samples with a wide range of amphiphilic protein concentrations. As expected, both amphiphilic proteins preferentially locate to the surface of condensates composed of RGG-RGG, within the range of concentrations tested (Figure 3D - E). With increasing concentrations of MBP-GFP-RGG we observe an increase in fluorescence at both the condensate interface and the aqueous phase, with little change in fluorescence inside condensates. For GST-GFP-RGG, an increase in protein concentration caused an increase in fluorescence signal mainly at the condensate interface, including phase separation of GST-GFP-RGG itself at the interface at the concentrations ≥ 5 µM, in agreement with our earlier study^19^.

We studied droplet sizes at the one-hour timepoint (Figure 3F - G). We observe a shift to smaller droplet sizes with increasing concentration for both amphiphilic proteins. Without addition of amphiphilic proteins, we observed a median droplet radius of 1.2 µm. Addition of high concentrations of either amphiphilic protein results in samples with condensates of a median radius of 0.5 – 0.6 µm. We fit our data to a sigmoidal equation:

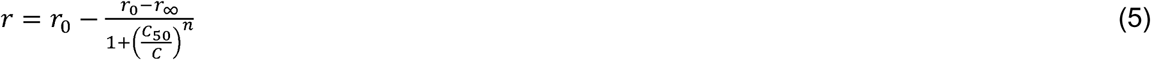

Here, *C* is the surfactant concentration, *r*_0_ and *r*_∞_ are the initial and final average droplet radii, *C*_50_ is the concentration required to reduce average condensate radius by half, and *n* is the Hill coefficient. We find that for MBP-GFP-RGG, *C*_50_ == 0.75 + 0.24 μM , and for GST-GFP-RGG, *C*_50_ == 0.18 + 0.08 μM . The difference in magnitude between these two fitting parameters is smaller than those measured for condensate interfacial tension. This is likely due to condensate size being regulated by additional factors beyond interfacial tension, for instance, surface charge may contribute to kinetic stability of condensates and other emulsions^35^. Notably, at high surfactant concentrations (> 5 µM), we observe droplets that remain in contact for long times without fusing, as mentioned previously (SI Appendix, Movie 1). In summary, our results suggest that the surfactant proteins that reduce interfacial tension also reduce droplet size.

## Conclusions

In this study, we quantify how engineered amphiphilic proteins modulate condensate interfacial tension. Our results directly demonstrate that the amphiphilic proteins act as surfactants and reduce the interfacial tension of condensates. We show that the degree of surfactant protein partitioning to the condensate surface determines its strength in lowering interfacial tension. By modulating the redox environment and thus preventing disulfide crosslinking between GST-GFP-RGG proteins, we can significantly reduce both the surfactant’s adsorption to the condensate interface and its capacity to lower interfacial tension. Furthermore, we also found that the reduction in interfacial tension results in slower fusion kinetics and reduced condensate sizes.

This work, together with other recent research^17,19^, suggests that protein-based surfactants may be one of several methods used to regulate the interfacial tension of condensates in cells. Given the importance of interfacial tension in determining the behavior of condensates^3^, these surfactant proteins may have extensive impacts on the function of condensates. For example, they may affect condensate multiphasic architecture, interaction with other intracellular structures, or size. Furthermore, we speculate that cells may control the strength of surfactants, such as by modulating the redox environment with the presence of reducing agents or using other mechanisms like posttranslational modification of proteins.

Looking ahead to bioengineering applications, amphiphilic proteins such as those described here can be used to improve engineered condensate systems. For example, engineered condensates have been explored as the basis for in vitro biocatalysis systems^36,37^. The ability to optimize biocatalysis in condensates by controlling condensate size, condensate surface area to volume ratio, and thus diffusion rates should be explored in the future.

## Acknowledgements

We thank Fleurie Kelley for helpful discussions. This work was supported by NIH grants R35GM142903 (to B.S.S.) and R35GM147027 (to Z.S), and National Science Foundation grant DMR-2238914 (to B.S.S.).

## Author Contributions

BF: conceptualization, experimental design, data collection and analysis, writing—original draft, writing—review and editing; HW: conceptualization, experimental design, data collection and analysis, writing—original draft, writing—review and editing; ZS: conceptualization, supervision, experimental design, data analysis, writing—original draft, writing—review and editing, funding acquisition; BSS: conceptualization, supervision, experimental design, data analysis, writing— original draft, writing—review and editing, funding acquisition.

## Resource Availability

Requests for further information and resources should be directed to and will be fulfilled by Benjamin S Schuster or Zheng Shi.

## Supporting Information

SI Appendix: Supplementary Notes containing protein sequences and equation derivations, Supplementary Table 1 and Supplementary Figures 1 – 4 with additional results and caption for Supplementary Video 1.

Supplementary Movie 1: additional fusion experiment results.

## References

1. Banani, S. F., Lee, H. O., Hyman, A. A. & Rosen, M. K. Biomolecular condensates: Organizers of cellular biochemistry. Nature Reviews Molecular Cell Biology vol. 18 285– 298 (2017).

2. Lyon, A. S., Peeples, W. B. & Rosen, M. K. A framework for understanding the functions of biomolecular condensates across scales. Nat. Rev. Mol. Cell Biol. 22, 215–235 (2021).

3. Wang, Z., Lou, J. & Zhang, H. Essence determines phenomenon: Assaying the material properties of biological condensates. Journal of Biological Chemistry vol. 298 (2022).

4. Guan, M., Hammer, D. A. & Good, M. C. Assembly of hierarchical multiphase condensates using designer surfactant proteins. *bioRxiv* (2024).

5. Feric, M. et al. Coexisting Liquid Phases Underlie Nucleolar Subcompartments Article Coexisting Liquid Phases Underlie Nucleolar Subcompartments. Cell 165, 1686–1697 (2016).

6. Quail, T. et al. Force generation by protein–DNA co-condensation. Nat. Phys. 17, 1007– 1012.

7. Agudo-Canalejo, J. et al. Wetting regulates autophagy of phase-separated compartments and the cytosol. Nature 591, 142–146 (2021).

8. Edward, G. & Shorter, J. The molecular language of membraneless organelles. J. Biol. Chem. 294, 7115–7127.

9. Gouveia, B. et al. Capillary forces generated by biomolecular condensates. Nature 609, 255–264 (2022).

10. Aarts, D. G. A. L., Schmidt, M. & Lekkerkerker, H. N. W. Direct Visual Observation of Thermal Capillary Waves. Science (80-. ). 304, 847–850 (2004).

11. Wang, H., Kelley, F. M., Milovanovic, D., Schuster, B. S. & Shi, Z. Surface tension and viscosity of protein condensates quantified by micropipette aspiration. Biophys. Reports 1, 100011 (2021).

12. Berry, J., Brangwynne, C. P. & Haataja, M. Physical principles of intracellular organization via active and passive phase transitions. *Reports Prog*. Phys. 81, 046601 (2018).

13. Meng, L., Mao, S. & Lin, J. Heterogeneous elasticity drives ripening and controls bursting kinetics of transcriptional condensates. Proc. Natl. Acad. Sci. 121, e2316610121 (2024).

14. Brangwynne, C. P., Mitchison, T. J. & Hyman, A. A. Active liquid-like behavior of nucleoli determines their size and shape in Xenopus laevis oocytes. Proc. Natl. Acad. Sci. U. S. A. 108, 4334–4339 (2011).

15. Folkmann, A. W., Putnam, A., Lee, C. F. & Seydoux, G. Regulation of biomolecular condensates by interfacial protein clusters. Science (80-. ). 373, 1218–1224 (2021).

16. Navarro, M. G-J. et al. RNA is a critical element for the sizing and the composition of phase-separated RNA–protein condensates. Nat. Commun. 10, 3230 (2019).

17. Wang, Z., Yang, C., Guan, D., Li, J. & Zhang, H. Cellular proteins act as surfactants to control the interfacial behavior and function of biological condensates. Dev. Cell 58, 919–932.e5.

18. Forman-Kay, J. D., Ditlev J. A., Nosella M. L., Lee H. O. What are the distinguishing features and size requirements of biomolecular condensates and their implications for RNA-containing condensates? RNA. 28, 36–47 (2022).

19. Kelley, F. M., Favetta, B., Regy, R. R., Mittal, J. & Schuster, B. S. Amphiphilic proteins coassemble into multiphasic condensates and act as biomolecular surfactants. Proc. Natl. Acad. Sci. 118, e2109967118 (2021).

20. Rosen, M. J. & Kunjappu, J. T. Surfactants and Interfacial Phenomena: Fourth Edition. Surfactants Interfacial Phenom. Fourth Ed. (2012) doi:10.1002/9781118228920.

21. Cuylen, S. et al. Ki-67 acts as a biological surfactant to disperse mitotic chromosomes. Nature 535, 308–312 (2016).

22. Wang, Z. et al. Material properties of phase-separated TFEB condensates regulate the autophagy-lysosome pathway. J. Cell Biol. 221, (2022).

23. Elbaum-Garfinkle, S. et al. The disordered P granule protein LAF-1 drives phase separation into droplets with tunable viscosity and dynamics. Proc. Natl. Acad. Sci. U. S. A. 112, 7189–7194 (2015).

24. Schuster, B. S. et al. Identifying sequence perturbations to an intrinsically disordered protein that determine its phase-separation behavior. Proc. Natl. Acad. Sci. U. S. A. 117, (2020).

25. Schuster, B. S. et al. Controllable protein phase separation and modular recruitment to form responsive membraneless organelles. Nat. Commun. 9, 2985 (2018).

26. Elbaum-Garfinkle, S. et al. The disordered P granule protein LAF-1 drives phase separation into droplets with tunable viscosity and dynamics. Proc. Natl. Acad. Sci. 112, 7189–7194 (2015).

27. Roggeveen, J. V., Wang, H., Shi, Z. & Stone, H. A. A calibration-free model of micropipette aspiration for measuring properties of protein condensates. Biophys. J. 123, 1393–1403 (2024).

28. Moazzeni, S. et al. N-Cadherin based adhesion and Rac1 activity regulate tension polarization in the actin cortex. Sci. Rep. 15, 4296 (2025).

29. Brugués, J. et al. Dynamical organization of the cytoskeletal cortex probed by micropipette aspiration. Proc. Natl. Acad. Sci. U. S. A. 107, 15415–15420 (2010).

30. Ghosh, A. & Zhou, H. X. Determinants for Fusion Speed of Biomolecular Droplets. Angew. Chemie - Int. Ed. 59, 20837–20840 (2020).

31. Goutelle, S. et al. The Hill equation: A review of its capabilities in pharmacological modelling. Fundam. Clin. Pharmacol. 22, 633–648 (2008).

32. Fainerman, V. B. et al. New view of the adsorption of surfactants at water/alkane interfaces – Competitive and cooperative effects of surfactant and alkane molecules. Adv. Colloid Interface Sci. 279, 102143 (2020).

33. Tudyka, T. & Skerra, A. Glutathione S_-_transferase can be used as a C_-_terminal, enzymatically active dimerization module for a recombinant protease inhibitor, and functionally secreted into the periplasm of Escherichia coli. Protein Sci. 6, 2180–2187.

34. Wang, H., Kelley, F. M., Milovanovic, D., Schuster, B. S. & Shi, Z. Surface tension and viscosity of protein condensates quantified by micropipette aspiration. Biophysical Reports vol. 1 (2021).

35. Welsh, T. J. et al. Surface Electrostatics Govern the Emulsion Stability of Biomolecular Condensates. Nano Lett. 22, 612–621 (2022).

36. Lim, D. & Clark, D. S. Phase-separated biomolecular condensates for biocatalysis. Trends Biotechnol. 42, 496–509.

37. Li, X. et al. Enzyme purification and sustained enzyme activity for pharmaceutical biocatalysis by fusion with phase_-_separating intrinsically disordered protein. Biotechnol. Bioeng. 121, 3155–3168.

