## Supplementary Notes 1 - 2, Supplementary Table, Figures S1 - S4, Caption SI Movie 1 for "Amphiphilic protein surfactants reduce the interfacial tension of biomolecular condensates"

**This PDF file includes:**

Supplementary Notes 1 - 2

Supplementary Table

Figures S1 - S4

Caption SI Movie 1

### **Supplementary Notes**

### **1. Protein sequences used in this work**

Throughout this paper, RGG denotes the RGG domain from LAF-1 (residues 1-168 of LAF-1). All proteins included a hexahistidine tag at the C-terminus for immobilized metal affinity chromatography. Below, RGG, GST, MBP, and GFP domains are color-coded; tags, linkers, and cut sites are not colored.

**RGG-RGG**

MESNQSNNGGSGNAALNRGGRYVPPHLRGGDGGAAAAASAGGDDRRGGAGGGGYRRGGGNSGGGGGGGYDRGYNDNRDDRDNRGGSGGYGRDRNYEDRGYNGGGGGGGNRGYNNNRGGGGGGYNRQDRGDGGSSNFSRGGYNNRDEGSDNRGSGRSYNNDRRDNGGDGEFGKLMESNQSNNGGSGNAALNRGGRYVPPHLRGGDGGAAAAASAGGDDRRGGAGGGGYRRGGGNSGGGGGGGYDRGYNDNRDDRDNRGGSGGYGRDRNYEDRGYNGGGGGGGNRGYNNNRGGGGGGYNRQDRGDGGSSNFSRGGYNNRDEGSDNRGSGRSYNNDRRDNGGDGLEHHHHHH

**MBP-GFP-RGG**

MKIEEGKLVIWINGDKGYNGLAEVGKKFEKDTGIKVTVEHPDKLEEKFPQVAATGDGPDIIFWAHDRFGGYAQSGLLAEITPDKAFQDKLYPFTWDAVRYNGKLIAYPIAVEALSLIYNKDLLPNPPKTWEEIPALDKELKAKGKSALMFNLQEPYFTWPLIAADGGYAFKYENGKYDIKDVGVDNAGAKAGLTFLVDLIKNKHMNADTDYSIAEAAFNKGETAMTINGPWAWSNIDTSKVNYGVTVLPTFKGQPSKPFVGVLSAGINAASPNKELAKEFLENYLLTDEGLEAVNKDKPLGAVALKSYEEELVKDPRIAATMENAQKGEIMPNIPQMSAFWYAVRTAVINAASGRQTVDEALKDAQTNSSSNNNNNNNNNNLGETVRFQSMVSKGEELFTGVVPILVELDGDVNGHKFSVSGEGEGDATYGKLTLKFICTTGKLPVPWPTLVTTLTYGVQCFSRYPDHMKQHDFFKSAMPEGYVQERTIFFKDDGNYKTRAEVKFEGDTLVNRIELKGIDFKEDGNILGHKLEYNYNSHNVYIMADKQKNGIKVNFKIRHNIEDGSVQLADHYQQNTPIGDGPVLLPDNHYLSTQSKLSKDPNEKRDHMVLLEFVTAAGITLGMDELYKGGGSENLYFQGEFGKLMESNQSNNGGSGNAALNRGGRYVPPHLRGGDGGAAAAASAGGDDRRGGAGGGGYRRGGGNSGGGGGGGYDRGYNDNRDDRDNRGGSGGYGRDRNYEDRGYNGGGGGGGNRGYNNNRGGGGGGYNRQDRGDGGSSNFSRGGYNNRDEGSDNRGSGRSYNNDRRDNGGDGLEHHHHHH

**GST-GFP-RGG**

MSPILGYWKIKGLVQPTRLLLEYLEEKYEEHLYERDEGDKWRNKKFELGLEFPNLPYYIDGDVKLTQSMAIIRYIADKHNMLGGCPKERAEISMLEGAVLDIRYGVSRIAYSKDFETLKVDFLSKLPEMLKMFEDRLCHKTYLNGDHVTHPDFMLYDALDVVLYMDPMCLDAFPKLVCFKKRIEAIPQIDKYLKSSKYIAWPLQGWQATFGGGDHPPGSETVRFQSMVSKGEELFTGVVPILVELDGDVNGHKFSVSGEGEGDATYGKLTLKFICTTGKLPVPWPTLVTTLTYGVQCFSRYPDHMKQHDFFKSAMPEGYVQERTIFFKDDGNYKTRAEVKFEGDTLVNRIELKGIDFKEDGNILGHKLEYNYNSHNVYIMADKQKNGIKVNFKIRHNIEDGSVQLADHYQQNTPIGDGPVLLPDNHYLSTQSKLSKDPNEKRDHMVLLEFVTAAGITLGMDELYKGGGSENLYFQGEFGKLMESNQSNNGGSGNAALNRGGRYVPPHLRGGDGGAAAAASAGGDDRRGGAGGGGYRRGGGNSGGGGGGGYDRGYNDNRDDRDNRGGSGGYGRDRNYEDRGYNGGGGGGGNRGYNNNRGGGGGGYNRQDRGDGGSSNFSRGGYNNRDEGSDNRGSGRSYNNDRRDNGGDGLEHHHHHH

### **2. Hill equation derivation**

Assuming that interfacial tension decreases linearly with the density of surfactant protein adsorbed to the interface of condensates, then:

$\gamma=\gamma_{0}-\alpha{\cdot\rho}_{s}$	(S1)

where *γ* is interfacial tension, *γ*_0_ is the interfacial tension without surfactant protein, *α* is a constant, and *ρ*_s_ is the density of surfactant protein adsorbed to the interface of condensates.

We propose that surfactant binding to the condensate interface can be modeled using a Hill equation:

$\rho_{s}=\frac{\rho_{\max}}{1+\left( \frac{K_{D}}{c} \right)^{p}}$	(S2)

where *ρ*_max_ is the maximum density of surfactant protein at the condensate interface, *c* is the surfactant concentration, and $K_{D}$ and *p* are the dissociation constant and cooperativity coefficient of the interfacial adsorption of the surfactants, respectively.

If we define $\gamma_{0}-\gamma_{\infty}=\alpha\rho_{\max}$, then:

$\gamma=\gamma_{0}-\frac{\gamma_{0}-\gamma_{\infty}}{1+\left( \frac{K_{D}}{c} \right)^{p}}$	(S3)

### **Supplementary Table**

Table 1: Inverse capillary velocities calculated from fusion data and micropipette aspiration data

| Protein Sample | Inverse Capillary Velocity from Fusion Experiments (s/µm) ± SEM | Inverse Capillary Velocity from Micropipette Aspiration Experiments (s/µm) ± SEM | Percent Difference  (%) |
| --- | --- | --- | --- |
| 10 µM RGG-RGG | 0.013 ± 0.001 | 0.014 ± 0.001 | 7.41 |
| 10 µM RGG-RGG +  1 µM MBP-GFP-RGG | 0.023 ± 0.004 | 0.025 ± 0.003 | 8.33 |
| 10 µM RGG-RGG +  5 µM MBP-GFP-RGG | 0.030 ± 0.004 | N/A | N/A |
| 10 µM RGG-RGG +  1 µM GST-GFP-RGG | 0.064 ± 0.005 | 0.047 ± 0.004 | 30.63 |
| 10 µM RGG-RGG +  5 µM GST-GFP-RGG | 0.104 ± 0.009 | N/A | N/A |

### **Supplementary Figures**

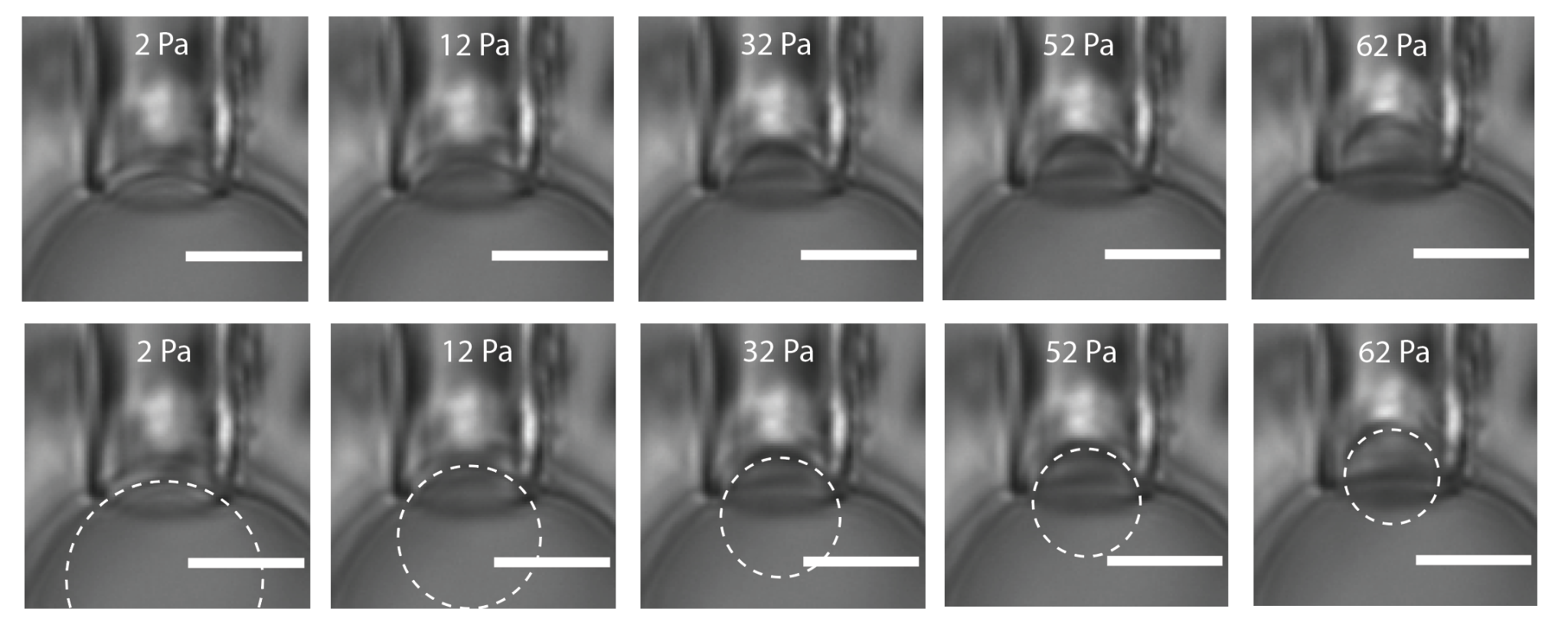

**Figure S1: Stepwise aspiration of an RGG-RGG condensate with 4 μM GST-GFP-RGG.** (Top) In 10 Pa increments, the condensate is gradually aspirated into the pipette until the deformation of the condensate interface is approximately equal to the inner radius of the micropipette. (Bottom) Dashed line indicates the fit to the radius of curvature of the deformed portion of the condensate.

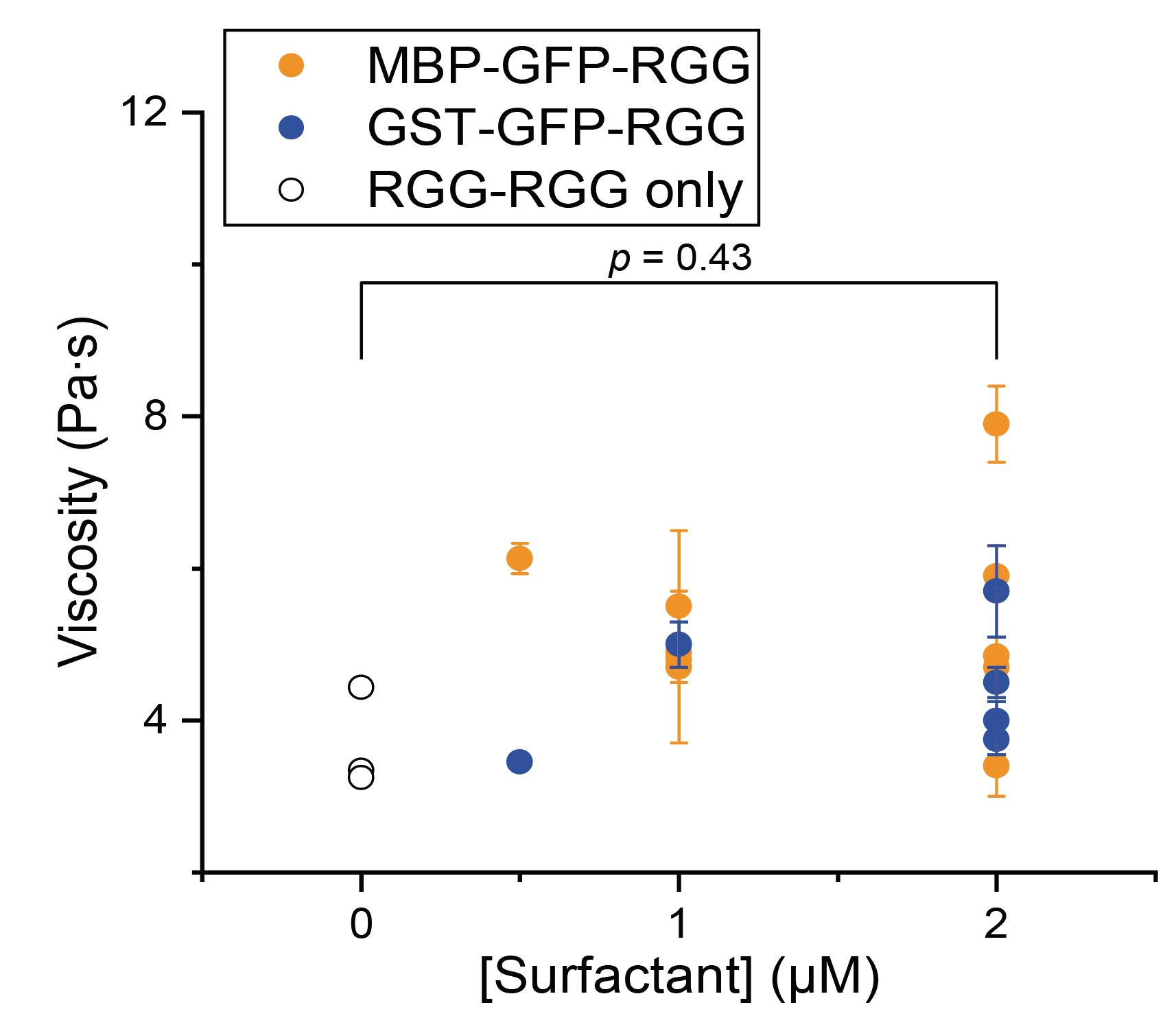

**Figure S2: Dependence of condensate viscosity on the concentration of surfactant protein added.** Each point shows the viscosity measured by micropipette aspiration for a single condensate, with error bars indicating the standard error for that measurement. Addition of surfactant does not significantly alter condensate viscosity; *p* values were determined using one-way ANOVA followed by post hoc Tukey’s test.

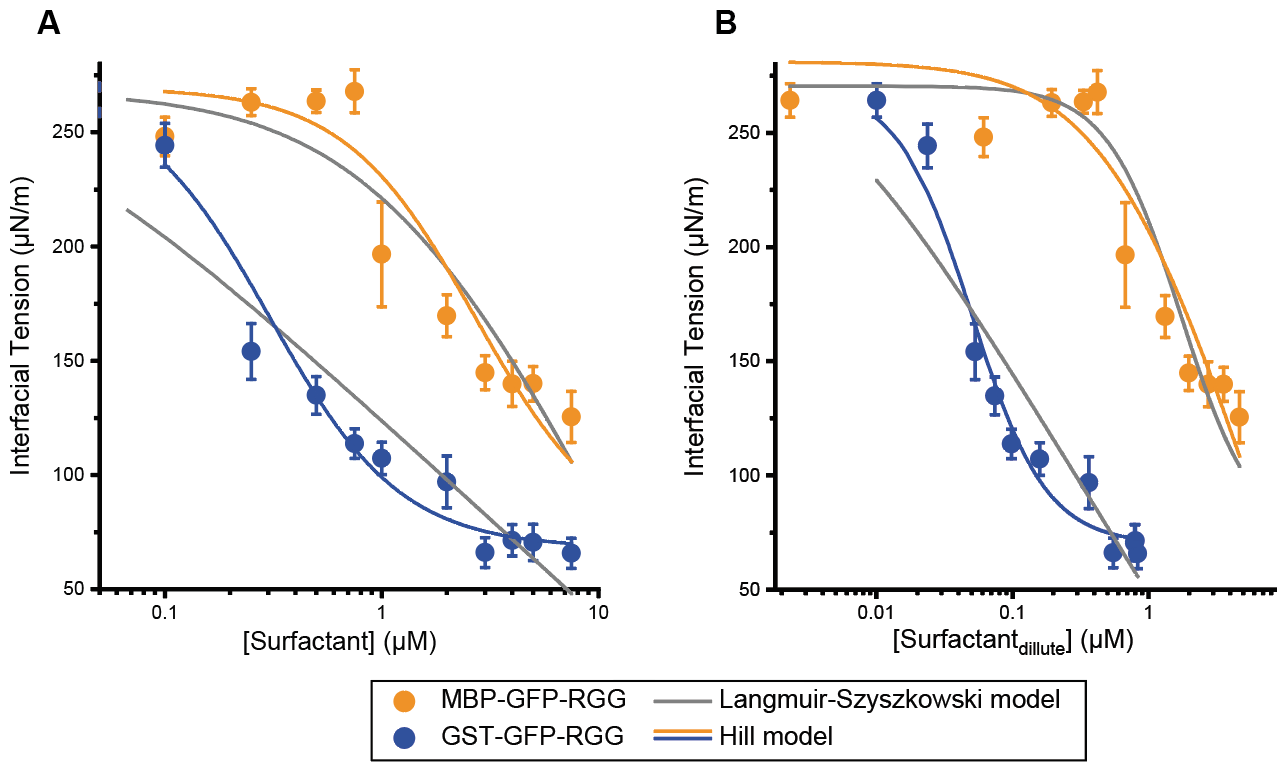

**Figure S3: Fitting of interfacial tension data with two adsorption models.** A) Interfacial tension of RGG-RGG condensates with increasing total concentrations of either amphiphilic protein. B) Interfacial tension of RGG-RGG condensates plotted against the equilibrium concentration of either amphiphilic protein in the dilute phase. The colored lines in panels A and B represent fitting curves using the Hill model (Eq. S3). Three fitting parameters ($\gamma_{0}$, $\gamma_{\infty}$ and *p*) are shared for the two amphiphilic proteins, while *K*_D_ is determined per protein (resulting in two parameters), meaning there are a total of 5 fitting parameters. The gray lines represent fitting curves from the Langmuir-Szyszkowski model: $\gamma=\gamma_{0}-B*\ln(1+KC)$, $B=RT\Gamma_{\infty}$ where $\gamma_{0}$ is the initial interfacial tension, *R* is the gas constant, *T* the absolute temperature, $\Gamma_{\infty}$ is the maximum possible adsorption at saturation, *C* is the surfactant concentration at equilibrium and $K$ is the adsorption equilibrium constant. The fitting parameter $\gamma_{0}$is shared for the two amphiphilic proteins, while *B* and *K* are determined per protein (resulting in four parameters), meaning that the fit also uses a total of 5 fitting parameters. Although both models have 5 fitting parameters, the R^2^ for the Hill model is 0.99 and 0.97 for panels A and B, respectively, while the R^2^ for the Langmuir-Szyszkowski model is 0.94 and 0.92 for panels A and B, respectively.

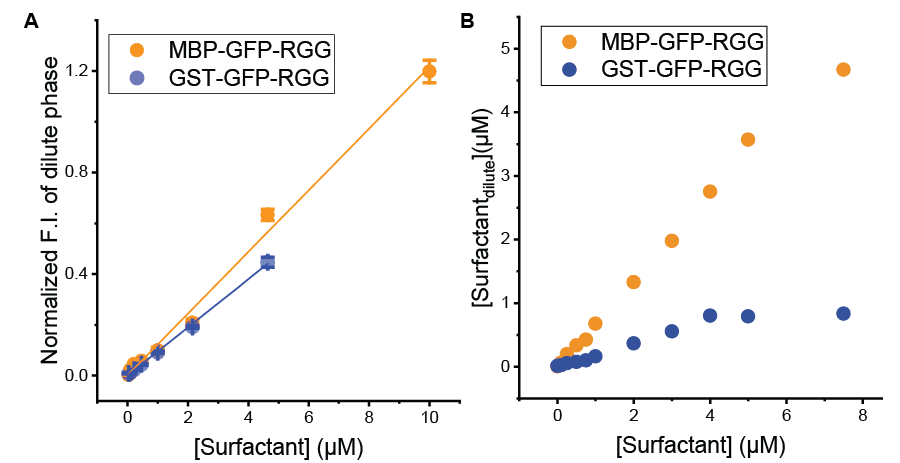

**Figure S4: Quantification of the amount of amphiphilic protein that remains in the dilute phase.** A) The calibration curves for MBP-GFP-RGG and GST-GFP-RGG. The relation between the normalized fluorescence intensity and the bulk surfactant concentration added to the sample (without RGG-RGG). Each point is the mean ± s.d. of five independent images; solid lines are linear fits used as calibration curves for converting fluorescence to absolute concentration (MBP-GFP-RGG: slope =${0.121\pm0.004 \mu M}^{-1}$, R^2^=0.98; GST-GFP-RGG: slope =${0.095\pm0.001 \mu M}^{-1}$, R^2^=0.99). B) Surfactant concentration in the dilute phase in samples with RGG-RGG, calculated using the equation from the calibration in A. MBP-based surfactant (orange) accumulates in the dilute phase more readily than the GST-based surfactant (blue) over the entire concentration range.

### **Supplementary Movies**

**SI Movie 1:** Optical tweezer-assisted droplet fusion experiment in a sample containing 10 µM RGG-RGG + 7.5 µM GST-GFP-RGG. Although droplets are approximated using optical tweezers, they do not fuse. Instead, we observe droplets moving vertically on top of each other.
